# Commensal skin bacteria exacerbate inflammation and delay skin healing

**DOI:** 10.1101/2023.12.04.569980

**Authors:** Veda D. Khadka, Laura Markey, Magalie Boucher, Tami D. Lieberman

## Abstract

The skin microbiome can both trigger beneficial immune stimulation and pose a potential infection threat. Previous studies have shown that colonization of mouse skin with the model human skin commensal *Staphylococcus epidermidis* is protective against subsequent excisional wound or pathogen challenge. However, less is known about concurrent skin damage and exposure to commensal microbes, despite growing interest in interventional probiotic therapy. Here, we address this open question by applying commensal skin bacteria at a high dose to abraded skin. While depletion of the skin microbiome via antibiotics delayed repair from damage, application of commensals-- including the mouse commensal *Staphylococcus xylosus*, three distinct isolates of *S. epidermidis,* and all other tested human skin commensals-- also significantly delayed barrier repair. Increased inflammation was observed within four hours of *S. epidermidis* exposure and persisted through day four, at which point the skin displayed a chronic-wound-like inflammatory state with increased neutrophil infiltration, increased fibroblast activity, and decreased monocyte differentiation. Transcriptomic analysis suggested that the prolonged upregulation of early canonical proliferative pathways inhibited the progression of barrier repair. These results highlight the nuanced role of members of the skin microbiome in modulating barrier integrity and indicate the need for caution in their development as probiotics.

## Introduction

In daily life, the gut, oral cavity, and skin are both a physical barrier to pathogens and a site of frequent immune crosstalk with resident microbes. These sites are subject to a daily barrage of superficial damage that allows for microbial entry across the physical barrier. Whether and how the host distinguishes friend from foe and how this impacts barrier health are foundational questions in understanding the host-microbe relationship. This work seeks to address how the host responds to commensal microbes when skin barrier integrity is compromised.

We focus on the skin because, among all barrier tissues, the skin provides a uniquely accessible and tractable system for the study of microbe-host interactions. Intact skin is composed of two layers: the outer epidermis, composed of proliferating keratinocytes and melanocytes protected by an outermost layer of cornified dead cells; and the inner dermis, composed of hair follicles, blood vessels, nerves and immune cells. The human skin microbiome is thought to colonize both the epidermal skin surface and hair follicles within the dermis (Conwill et al. 2022; Acosta et al. 2023).

In adulthood, the healthy human skin microbiome is dominated by *Cutibacterium acnes, Staphylococcus epidermidis* and *Corynebacterium* species (Grice et al. 2009). These commensal microbes are thought to protect the host against opportunistic pathogens by competing for resources (Wei et al. 2023), engaging in microbial warfare via the secretion of antimicrobial peptides (Nakatsuji et al. 2017; Wei et al. 2023), and interfering with pathogen quorum sensing (Williams et al. 5 2019; Canovas et al. 2016; Ramsey et al. 2016; Paharik et al. 2017). When the integrity of the skin barrier is compromised, as occurs in atopic dermatitis patients during disease flares, opportunistic pathogens such as *Staphylococcus aureus* can exploit the damaged barrier, dominate the skin microbial community, and secrete cytotoxic factors that trap the host in a chronic inflammatory loop (Kong et al. 2012), (Wanke et al. 2013; Nakagawa et al. 2017). The inability of these opportunistic pathogens to cause damage when the barrier is intact highlights the role of host barrier integrity in determining the response to pathogen exposure.

Comparatively little is known about how the host responds to commensal microbes when barrier integrity is compromised. Previous work in this area has primarily used full-thickness excisional wound and infection models, in which all layers of the skin are removed or damaged. In these models, colonization is beneficial: germ-free mice mount impaired immune responses (Uberoi et al. 8 2021; Naik et al. 2012; Wang et al. 2021), and conventional mice colonized with *S. epidermidis* prior to damage display improved healing outcomes (Uberoi et al. 8 2021; Naik et al. 2012; Wang et al. 2021). A few studies have used simultaneous damage and *S. epidermidis* exposure but have observed contradictory responses. Following mild epidermal abrasion, *S. epidermidis* application promoted skin repair (Zheng et al. 2022). However, in a mouse model of atopic dermatitis (Cau et al. 2021), and after strong tape-stripping and *S. aureus* co-infection (Burian et al. 2017), *S. epidermidis* application increased pathology and disrupted skin barrier repair.

Here, we use moderate tape-stripping to disrupt the upper layers of the epidermis and apply commensal bacteria daily to the damaged skin. Unlike previous studies, we apply a high microbial load throughout the period of study to model topical probiotic treatment of damaged skin and test the effects of a range of bacterial species, from pathogens to well-characterized probiotics. We find that application of all tested opportunistic pathogens and commensal skin bacteria, including three isolates of the model commensal *S. epidermidis,* delayed healing from epidermal abrasion. Delayed healing in response to *S. epidermidis* was mediated by amplification of the innate immune response to damage, expression of the cytokine IL-17A, and aberrantly prolonged epithelial proliferation. Our results suggest that application of most skin commensal bacteria to a damaged skin barrier can be detrimental to the host and may be unsuitable for development into therapeutic applications.

## Results

### Perturbation of the skin microbiome affects recovery from barrier repair

To investigate how native skin commensals affect barrier repair from epidermal abrasion, we first disrupted the skin barrier by tape-stripping (Fig. 1a). Hair was removed from the back of each mouse and Tegaderm (3M) was repeatedly applied and removed in order to abrade much of the epidermis (Fig. 1a and Fig. S1f). Histopathological analysis showed that tape-stripping resulted in erosion of the top layers of the epidermis and rarely extended to the dermis, indicating that abrasion by tape-stripping is distinct from full-thickness wounding (Fig. S1f). Each day, 100 μl of washed bacterial cells or phosphate-buffered saline vehicle (PBS) was pipetted onto damaged skin and gently spread across the skin surface using a sterile cotton swab. Application occurred immediately after tape-stripping and then daily until experimental endpoint. Barrier recovery was assessed daily by transepidermal water loss (TEWL) and a severity score that semi-quantitatively assessed erythema (redness), crust, and thickness (Fig. S1c-d). To quantify healing over time and enable comparison between cohorts, the TEWL and severity score trajectories for each animal were summarized using area under the curve (AUC) and normalized to the average values for the control group from that cohort (Fig. S2a-b).

**Figure 1:**
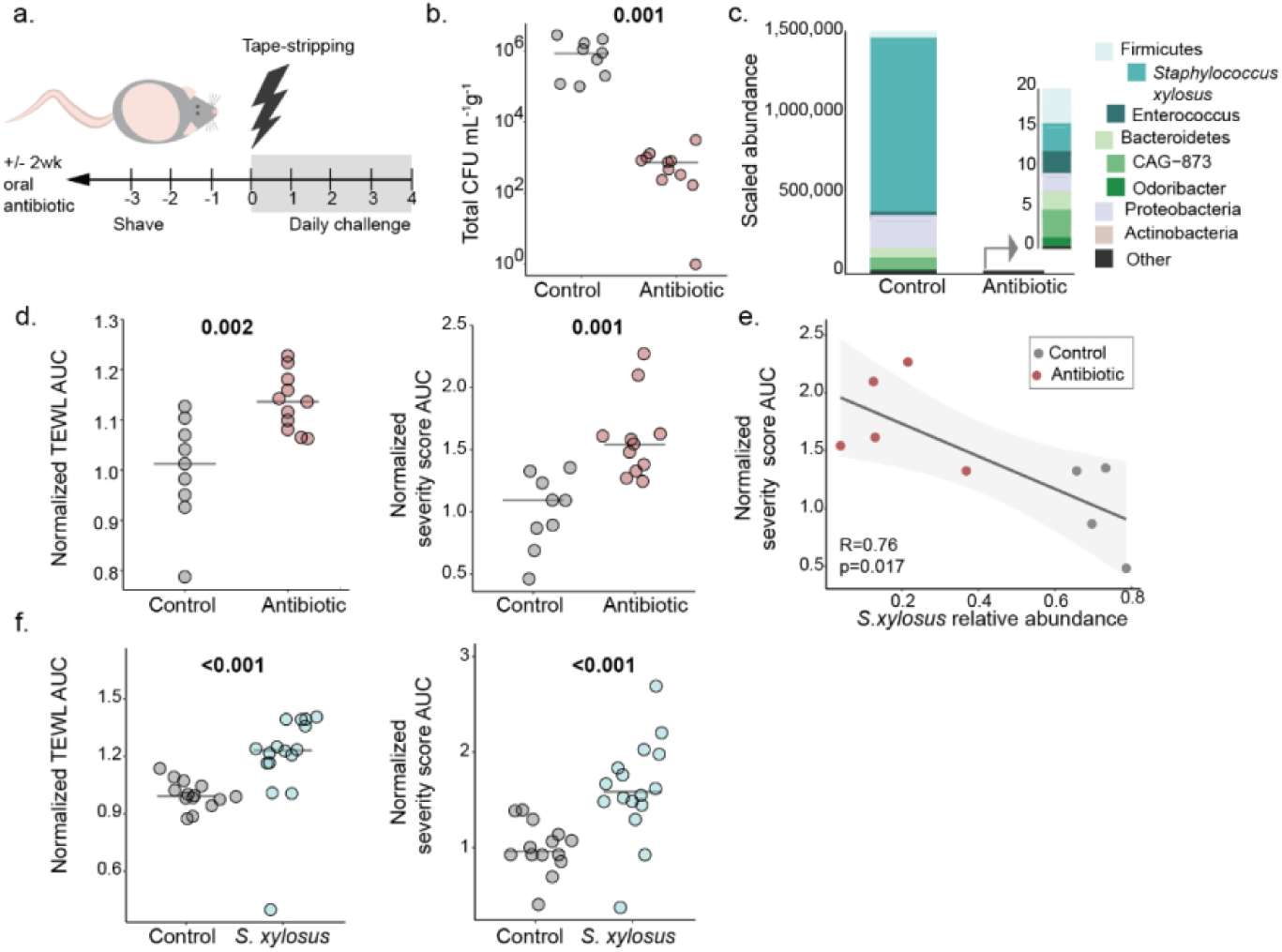
Perturbation of the native skin microbiome by depletion or supplementation delays healing. a) Mice received antibiotics for two weeks prior to hair removal and tapestripping followed by daily application of either PBS (control) or bacteria of interest. b) Skin microbiota endpoint viable bacteria measured from skin homogenate plated on mannitol salt agar (control N=9, antibiotics N=11). c) 16s amplicon sequencing of endpoint skin swabs (control N=4, antibiotics N=5). Scaled abundance=average CFUs*average taxon abundance. d) Skin barrier integrity assessed by TEWL (left) and severity score (right) which were summarized by area under the curve (AUC) and normalized to cohort control average (control N=9, antibiotics N=11). e) Correlation between *S. xylosus* relative abundance and TEWL AUC (top) and severity score AUC (bottom). f) TEWL (left) and severity score (right) measured daily and summarized by normalized AUC (control N=10, *S. xylosus* N=10). For strip-plots, symbols represent mice and bars indicate medians.

To determine the role of the native skin microbiome in repair from abrasion, we depleted the microbiome of Specific Pathogen Free (SPF) mice, using an antibiotic cocktail designed to target the skin flora (Uberoi et al. 8 2021). We confirmed that this treatment depleted recoverable bacteria on the skin ∼10,000-fold (Fig. 1b-c, P=0.001), while only minimally impacting the gut microbiome (Fig. S2g-i).

Antibiotic-treated mice had higher TEWL and severity scores compared to control mice for the entirety of the exposure period (Fig. 1d, Fig. S2a-b, P<0.003), suggesting that depletion of the skin microbiome delayed healing. Control mice (which healed faster) had a less diverse skin microbiome when compared to antibiotic-treated mice at the end of the exposure period (Fig. S2c-f, P=0.016). This result was surprising given many examples of low diversity communities being associated with disease severity (Kong et al. 2012; Gevers et al. 2014; Clausen et al. 2018). *Staphylococcus xylosus,* a species commonly recovered from mouse skin during health ((Belheouane et al. 2020), dominated the skin microbiomes of these control mice (Fig. 1c and Fig. S2c-d). In line with a beneficial role for *S. xylosus* after tape-stripping, the endpoint relative abundance of *S. xylosus* was negatively correlated with skin damage as assessed by severity score AUC (Fig. 1d, R=-0.76, P=0.017) or TEWL AUC (Fig. S2a, R=-0.53, P=0.14) across both control and antibiotic-treated mice. These results suggest that a decreased abundance of native *S. xylosus* could delay barrier recovery.

We therefore hypothesized that supplementation of the skin microbiome with exogenous *S. xylosus* might improve healing in mice with a complete microbiome. To test this hypothesis, we cultured commensal *S. xylosus* from SPF mice and applied either PBS (vehicle control) or 10^9^ CFUs immediately following tape-stripping damage and daily until experimental endpoint. Contrary to our hypothesis and in line with previous reports of *S. xylosus*-exacerbated disease in compromised skin (Gimblet et al. 7 2017; Won et al. 2002; Reshamwala et al. 2022), mice exposed to *S. xylosus* displayed delayed healing by both elevated TEWL and severity score compared to control mice (Fig. 1f, P<0.001). Thus, either depletion or addition of the native mouse commensal *S. xylosus* after barrier damage delays healing.

### Additional commensal skin bacteria do not improve skin healing following mechanical damage

We next tested if the delayed recovery from barrier damage was specific to supplementation with *S. xylosus* or generalizable across diverse skin commensals. To test this, we selected a variety of members of the healthy human skin microbiome previously shown to improve barrier immunity (Naik et al. 2015; Nakatsuji et al. 2021; Zheng et al. 2022): *S. epidermidis*, *Staphylococcus hominis*, and *Corynebacterium accolens*, as well as a community-associated, methicillin-resistant isolate of *S. aureus (*USA300). For all species, exponential phase bacterial cultures were washed in PBS and 10^8^-10^9^ CFUs were applied to damaged skin on a daily basis throughout the experiment.

Surprisingly, all tested human skin commensal microbes significantly delayed healing relative to controls, by both TEWL and severity score (Fig. 2a, P<0.03). Application of the opportunistic pathogen *S. aureus* also significantly delayed healing by both measures (Fig. 2a, P<0.001). Mice exposed to *S. aureus* were substantially more damaged than those exposed to commensals, displaying the highest severity score possible (Fig. S3), in line with an expected difference in immune response between commensals and pathogens.

**Figure 2:**
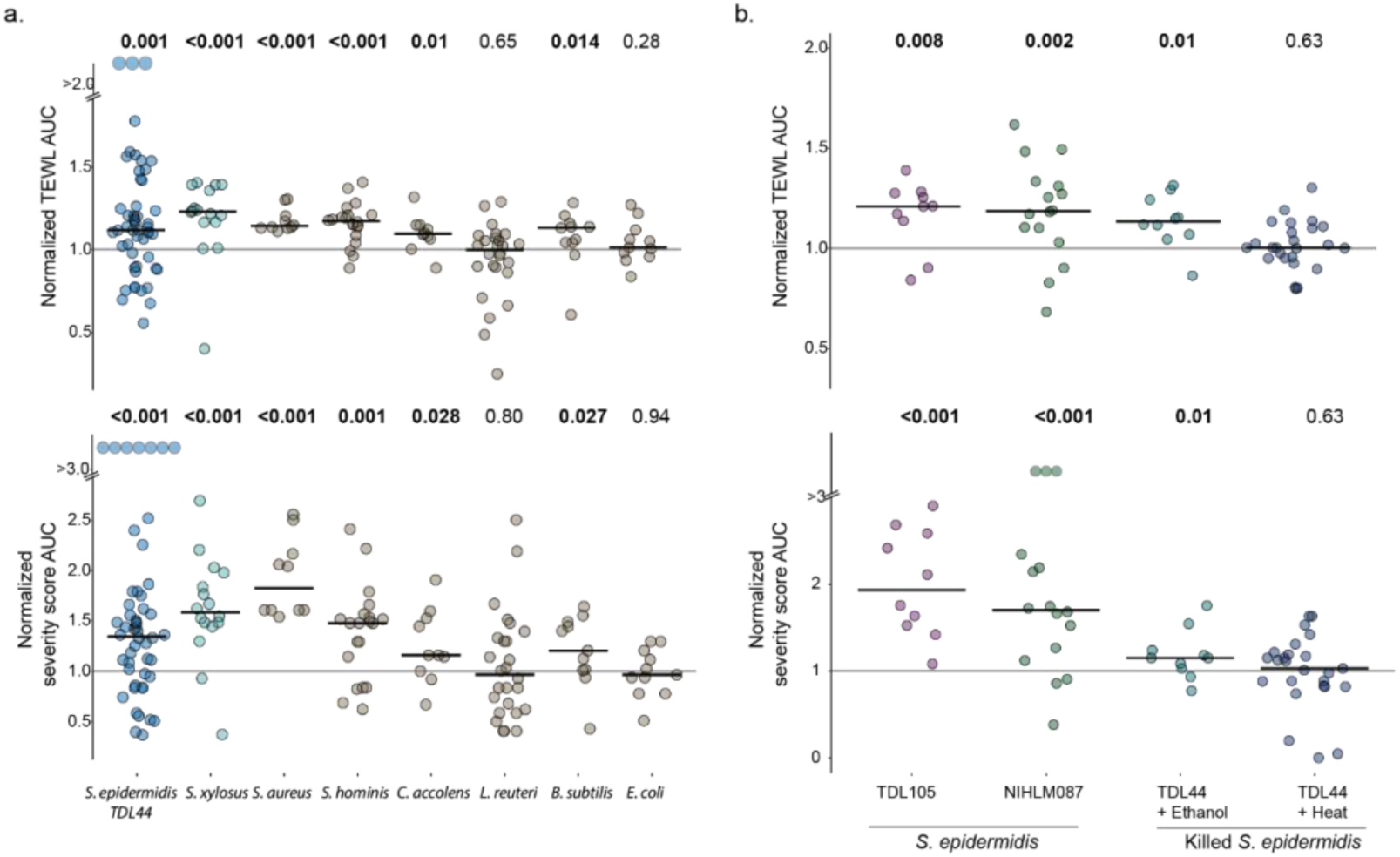
Human skin commensals including *S. epidermidis* delay barrier repair. a) PBS (control) or listed bacteria was applied to skin after tapestripping and then daily until endpoint. TEWL and severity score trajectories for each animal were summarized using area under the curve (AUC) and normalized to the cohort control average (represented as y=1). TEWL (top) and severity score (bottom) AUCs were compared to control average. b) TEWL (top) and severity score (bottom) AUCs after application of two additional isolates of *S. epidermidis* (left) or heat-killed or ethanol-killed Sepi-TDL44 (right). Throughout, symbols indicate individual mice and bars represent medians. Sepi-TDL44 N=51, *S. xylosus* N=16, *S. aureus* N=10, *S hominis* N=20, *C. accolens* N=10, *L. reuteri* N=27, *B. subtilis* N=11, *E. coli* N=11, Sepi-TDL105 N=10, Sepi-NIHLM087 N=16, ethanol-killed Sepi-TDL44 N=10, heat-killed Sepi-TDL44 N=20. Results typically include 2-3 cohorts per experimental condition.

We next tested if this trend of delayed skin healing was a universal response to microbial exposure during damage. We tested laboratory isolates of *Escherichia coli* (MG1655) and the soil bacterium *Bacillus subtilis*, as well as a human isolate of the putative gut probiotic *Limosilactobacillus reuteri* (ATCC 23272) in our tape-stripping model. Neither *E. coli* nor *L. reuteri* delayed healing over controls when applied to tape-stripped skin (Fig. 2a, P>0.3). In contrast, mice challenged with *B. subtilis* during damage displayed significantly elevated TEWL and severity scores (Fig. 2a, P<0.03). *B. subtilis* has been found in infected burn wounds (Saleh et al. 2014) and is sometimes considered an opportunistic pathogen, thus its deleterious effect on skin healing is not completely unexpected. These results show that the detrimental effect of microbial exposure on barrier repair is not common to all bacteria applied to damaged skin and appears to be limited to skin commensals and opportunistic pathogens.

### Multiple isolates of S. epidermidis delay healing when applied during damage

Given prior work on the beneficial properties of *S. epidermidis* in mouse models, we sought to ascertain whether the observed delay in healing was unique to the tested isolate (Sepi-TDL44, isolated from healthy volunteers) or a generalized feature of *S. epidermidis.* We tested a *S. epidermidis* isolate previously characterized as beneficial in wound healing, Sepi-NIHLM087 (Linehan et al. 2018), as well as a second commensal isolate of *S. epidermidis* from healthy volunteers (Sepi-TDL105). Both Sepi-NIHLM087 and Sepi-TDL105 delayed healing significantly over controls similarly to Sepi-TDL44 (Fig. 2b, P<=0.008), and there was no significant difference between isolates (P>0.5). These data confirm that delayed skin healing is a generalized host response to *S. epidermidis* and not a strain-dependent effect.

In contrast to the delayed healing observed when Sepi-TDL44 was applied to damaged skin, daily application to depilated healthy skin did not induce any morphological changes or impact skin health as measured by TEWL or severity score (Fig. S4a-b). Moreover, *S. epidermidis* exposure during health did not change the response to subsequent barrier damage (Fig S4c-d). This neutral response to Sepi-TDL44 applied to healthy skin demonstrates a clear difference in the host response to Sepi-TDL44 depending on skin barrier health during exposure.

To test whether live bacterial activity was required for delayed healing, mice were exposed to either heat-killed Sepi-TDL44 or ethanol-killed Sepi-TDL44. Application of heat-killed bacteria (boiled lysates of washed cells at equivalent cell density) did not delay skin healing (Fig. 2b, P=0.63). In contrast, when bacteria were exposed to 80% ethanol (which kills bacteria through coagulation and disruption of their cell membranes (Taddese et al. 2021)), washed, and applied to damaged skin, healing was significantly delayed over controls (Fig. 2b, P=0.01) - indicating that the delayed healing response was not solely dependent on secreted products. These results agree with reports that the interaction of tape-stripped skin with a cell-bound component of *S. epidermidis* worsens disease (Burian et al. 2017).

### Application of S. epidermidis after damage induces an inflammatory innate response

To better understand how *S. epidermidis* changes the host response to damage, we conducted histopathological analysis at multiple timepoints after tape-stripping. By pathology score (Methods), tape-stripped skin exposed to Sepi-TDL44 was less healed four days after tape-stripping damage (Fig. 3a and Fig. S1e), in line with TEWL and severity score measurements. Sepi-TDL44-exposed mice displayed marked epidermal ulceration with serocellular crust, as well as inflammatory infiltrates consisting primarily of neutrophils, fibroblasts, and some mast cells (Fig. 3a). In contrast, controls displayed lesions that were much more healed, characterized by epidermal hyperplasia, hyperkeratosis, and increased dermal fibroblasts.

**Figure 3:**
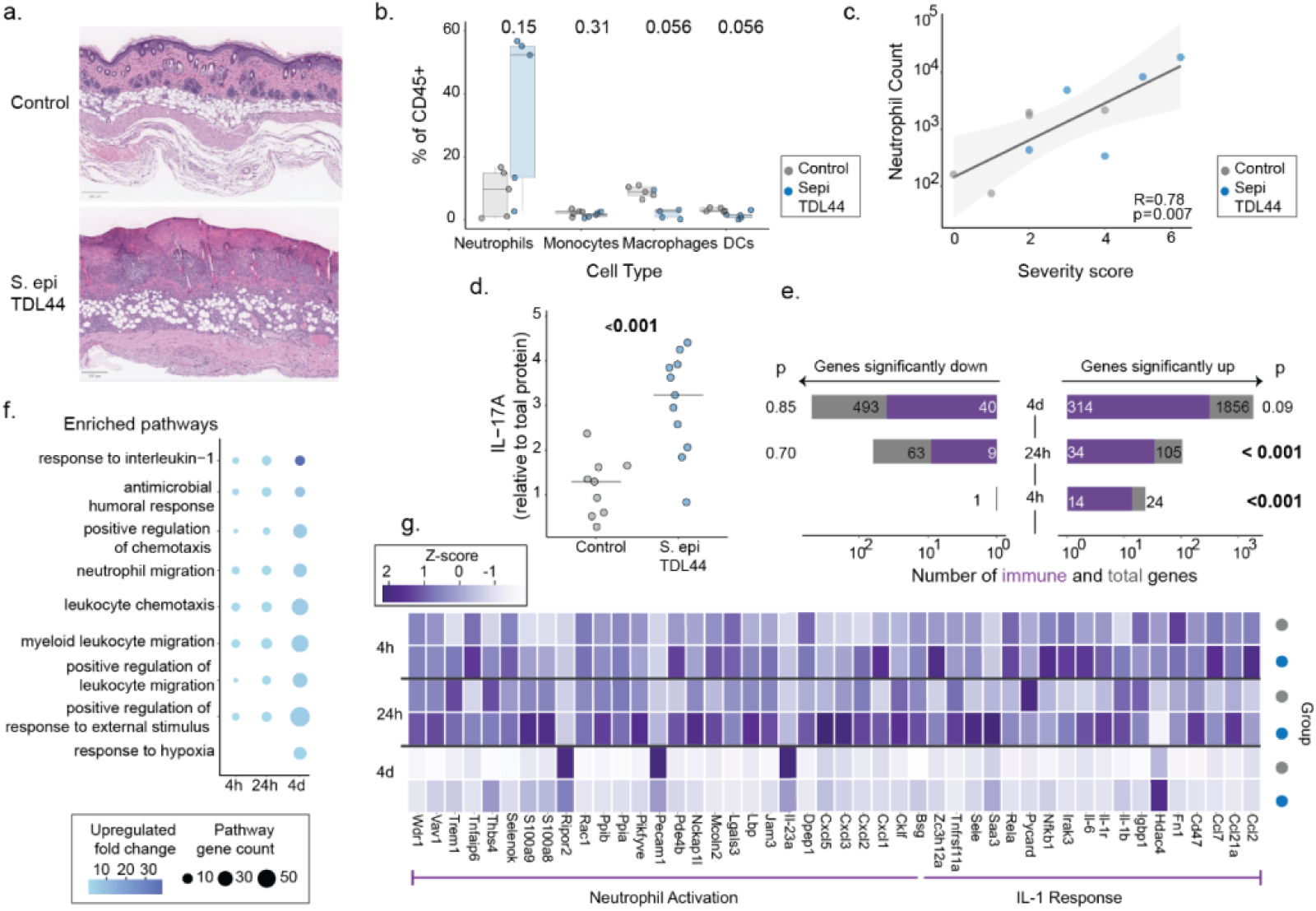
Application of *S. epidermidis* following barrier damage induces an innate inflammatory immune response. PBS (control) or Sepi-TDL44 was applied to skin after tapestripping and then daily until endpoint. a) Representative H&E-stained flank skin four days after damage. b) Flow cytometry performed four days after damage and PBS/Sepi-TDL44 application (control N=5, Sepi-TDL44 N=5). c) Correlation between neutrophils and day 4 raw severity score. d) IL-17A protein measured in skin lysate from skin collected three days after damage (control N=9, Sepi-TDL44 N=11). e) Bulk skin RNAseq differential expression analysis using likelihood ratio test to identify significantly differential genes (total height of gray bar) in response to Sepi-TDL44 and immune gene enrichment (purple bar) (4h control N=5, Sepi*-*TDL44 N=5; 24h control N=4, Sepi*-*TDL44 N=5; 4 days control N=6, Sepi*-*TDL44 N=9). f) Pathway enrichment analysis of Sepi-TDL44-upregulated immune genes. g) Average normalized gene expression (TPM). Symbols indicate individual mice and bars indicate median.

Next, we wanted to define immune pathways contributing to the delayed healing observed in response to *S. epidermidis*. Previous reports have shown the adaptive immune response to application of Sepi*-*NIHLM087 on healthy skin (induction of IL-17A producing CD8+ T-cells 14 days after bacterial application) is protective against pathogen challenge and excisional wounding (Naik et al. 2015). On day seven post-damage and Sepi-TDL44 exposure, we measured an increase in both bulk T-cells and γδ-T cells specifically (Fig. S5c, P=0.004 and P=0.0087 respectively) in response to Sepi-TDL44; however, skin barrier repair remained delayed relative to controls (Fig. S5a-b). This finding suggests that the T cell response to *S. epidermidis* does not promote healing from tape-stripping.

To identify additional immune pathways involved in the delayed healing response to *S. epidermidis*, we used a multiplex ELISA to characterize the cytokines (Fig. S6) produced in the skin three days post-damage. We measured a significant increase only in IL-17A in mice exposed to Sepi-TDL44 (Fig. 3d, P<0.001; p-value threshold corrected for multiple hypotheses).

Previous work has shown that skin IL-17A can have many sources including CD8+ (Naik et al. 2015) and CD4+ T cells (Asarch et al. 2008), γδ-T cells (Cai et al. 2011), and innate immune cells including neutrophils and mast cells (Lin et al. 2011). Given the rapidity of the inflammatory response observed after Sepi-TDL44 exposure, we next used flow cytometry to quantify innate immune cell populations in the skin.

We performed flow cytometric analysis four days after concurrent damage and *S. epidermidis* exposure to capture innate immune cell populations present. We observed an increase in neutrophils in mice exposed to Sepi-TDL44 compared to controls, but a decrease in monocytes, macrophages, and dendritic cells (Fig. 3B). Supporting a pathological role for excess neutrophil recruitment, mouse skin severity scores were strongly and significantly correlated with the number of neutrophils (Fig. 3c, R=0.78, P=0.007). Altogether, our flow cytometry results indicate that *S. epidermidis* induces both innate and adaptive immune cell populations, which could interfere with the precise succession of immune cells and cytokines required for successful initiation and resolution of the first, inflammatory, phase of skin wound healing (Landén et al. 2016).

Transcriptomic analysis of the damaged flank skin from three acute timepoints after damage (4h, 24h and 4 days) confirmed strong induction of the innate immune response in mice exposed to Sepi-TDL44. Significantly upregulated genes were enriched in gene annotations related to the immune system in Sepi-TDL44-exposed mice at 4h and 24h after damage (Fig. 3e, P<0.001). More specifically, the IL-1 response pathway was upregulated in Sepi-TDL44-exposed mice at all timepoints, in agreement with multiple literature reports of IL-1 activation after *Staphylococcus sp*. exposure on the skin (Leech et al. 2019; Wang et al. 2021). Additionally, immune cell chemotaxis pathways were enriched among upregulated genes at all timepoints (Fig. 3f and Table S1), including pathways for neutrophil migration (consistent with histopathology and flow cytometry, Fig. 3a-b), leukocyte migration, and general chemotaxis.

Many of the immune pathways upregulated by Sepi-TDL44 are part of the normal inflammatory phase of wound healing, which typically lasts 0-3 days (Singh et al. 2017). Critically, however, expression of genes in these pathways decreased over time in controls but remained higher in Sepi-TDL44-exposed mice (Fig. 4c, Fig. S7). This led to their relative upregulation compared to controls at later timepoints. Thus, Sepi-TDL44 stimulation exacerbated normal inflammation and extended it beyond its expected duration of normal wound healing.

**Figure 4:**
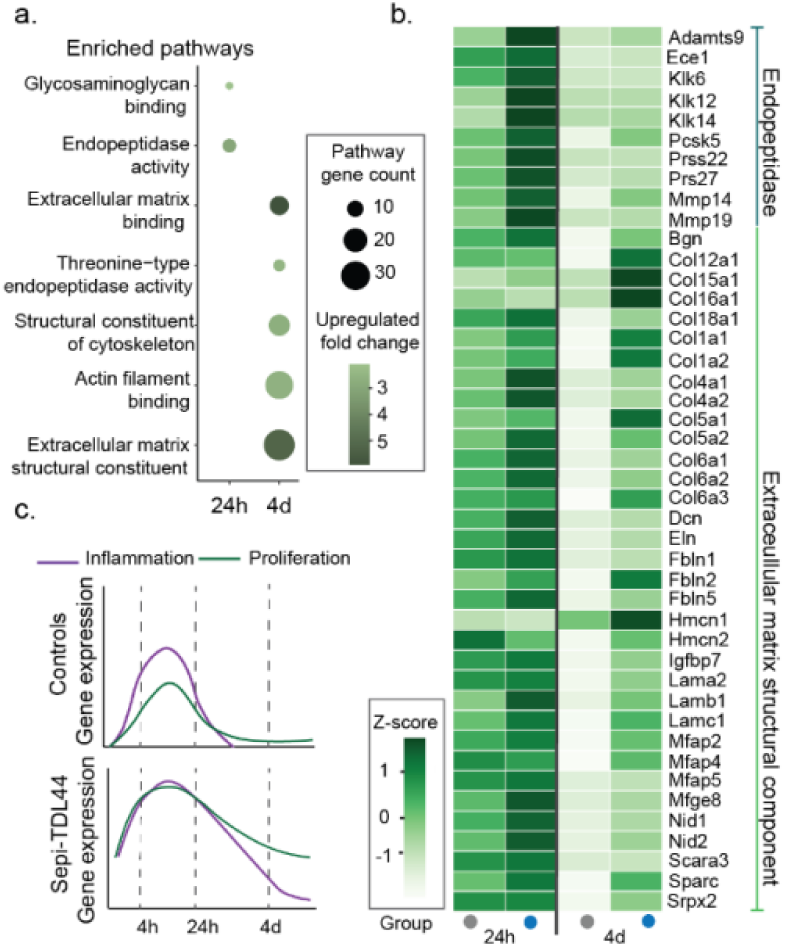
*S. epidermidis* application after barrier damage increases and prolongs expression of epithelial cell proliferation pathways. PBS (control) or Sepi-TDL44 was applied to skin after tapestripping and then daily until endpoint (4h, 24h or 4 days). a) Pathway enrichment analysis of non-immune genes upregulated in Sepi-TDL44-exposed mice relative to controls. b) Average normalized gene expression (TPM) of selected GO processes shown in c (4h control N=5, Sepi*-*TDL44 N=5; 24h control N=4, Sepi*-*TDL44 N=5; 4 days control N=6, Sepi*-*TDL44 N=9). c) Cartoon illustrating that mice exposed to Sepi-TDL44 after damage show an increased and prolonged expression of inflammation and proliferation genes compared to controls. For all RNAseq analysis, gene-level statistical significance comparing Sepi-TDL44-exposed mice to controls was calculated using the likelihood ratio test with Benjamini-Hochberg correction for multiple comparisons.

### S. epidermidis prolongs expression of epithelial proliferation pathways

In healing wounds, immune cell recruitment is followed by re-epithelization through cell migration and proliferation. Inflammatory cytokines such as IL-1 stimulate epithelial cell proliferation, and products of proliferation such as extracellular matrix (ECM) fragments further stimulate inflammation (Adair-Kirk and Senior 2008). This creates a positive feedback loop that promotes re-epithelization and is normally terminated upon production of anti-inflammatory factors with healing progression (Michopoulou and Rousselle 2015). In addition to amplifying the innate immune response, *S. epidermidis* exposure also upregulated pathways involved in these downstream proliferative phases of healing. Sepi-TDL44-exposed mice showed prolonged expression of genes involved in ECM and epithelial proliferation, with significantly increased expression at day 4, indicating aberrant elongation of this phase of healing (Fig. 4 and Table S2).

Among re-epithelization pathways upregulated by Sepi-TDL44, host-produced protease expression was particularly noteworthy. Multiple classes of ECM-targeting endopeptidases were upregulated 24h after damage, including kallikrein, matrix metalloprotease, ADAM metalloproteinase and threonine-type protease gene families (Fig. 4a-b). While protease activity is required to debride the damaged skin and provide access for migrating cells (Hattori et al. 2009; Michopoulou and Rousselle 2015), persistent protease activity produces proinflammatory ECM fragments (Adair-Kirk and Senior 2008) and degrades factors required for skin closure (Herrick et al. 1997; Lauer et al. 2000). Excessive protease expression has been shown to directly delay healing in elderly patients (Herrick et al. 1997). Sepi-TDL44-exposure also upregulated skin and ECM structural proteins (Fig. 4a-b), including glycosaminoglycan binding proteins, collagen, and other ECM/cytoskeleton structural constituents. This upregulation suggests increased influx and activity of fibroblasts (Bergmeier et al. 2018). Consistent with this result, histopathology showed increased fibroblasts within the dermis of Sepi-TDL44-exposed mice at sites of delayed healing (Fig. 3a). Together these results indicate that delayed healing in the skin of Sepi-TDL44-exposed mice is driven by an excess of innate inflammation and prolonged cellular proliferation (Fig. 4c).

## Discussion

In health, the skin microbiome can play a protective role by promoting barrier integrity and providing colonization resistance against opportunistic pathogens (Wei et al. 2023). Our results show that, despite their beneficial role during health, the application of native human and mouse skin commensals to damaged skin delays healing by exacerbating inflammation and prolonging proliferation. Our results also highlight the importance of host barrier context in influencing the response to commensal microbes: both depletion and supplementation of the native skin microbiome during damage are harmful for barrier repair, though others have shown responses may be beneficial during health.

Interestingly, our results indicate that IL-17A induction by *S. epidermidis* in the context of barrier damage delays healing, while induction of the same cytokine by *S. epidermidis* applied to healthy skin has been shown to boost beneficial T-cell responses, which defend against infection and promote wound closure (Naik et al. 2012, 2015; Linehan et al. 2018). Thus, host barrier context plays a critical role in determining the consequence of cytokine induction by a commensal. The increased IL-17A expression in *S. epidermidis*-exposed mice with delayed barrier repair is reminiscent of diabetic and infected wounds, where IL-17A expression has been implicated in delayed healing (Lee et al. 2018; Lecron et al. 2022). Similarly, the innate cell imbalance observed in *S. epidermidis*-exposed mice, created by a prolonged influx of neutrophils and absence of macrophages, is also present in chronic wounds (Joshi et al. 9 2020). Supporting a causal role for IL-17A in delaying repair, mice deficient in IL-17A have fewer neutrophils and accelerated wound closure (Takagi et al. 2 2017). Our results add to the body of work demonstrating a pathological role for excessive innate inflammation in skin healing.

Our results suggest that commensal bacteria could amplify inflammation and other disease pathology in people with a weakened or inflamed skin barrier, such as patients with atopic dermatitis. Shifts in microbiome composition are common among atopic dermatitis patients, and Staphylococci (largely *S. aureus* but occasionally *S. epidermidis* and other coagulase-negative Staphylococci) generally dominate the inflamed skin microbiome (Byrd et al. 2017; Chang et al. 2018; Williams et al. 2020; Khadka et al. 2021). Increased endogenous proteases are also associated with pathology in atopic dermatitis patients (Komatsu et al. 2007; Voegeli et al. 2009; Nomura et al. 2020), echoing our results of Staphylococci-induced upregulation of host protease production. While future work will be needed to understand the clinical implications of our work given the structural and regenerative differences between mouse and human skin (Zomer and Trentin 2018), our results highlight several pathways via which *S. epidermidis*, and potentially other commensal skin bacteria, can exacerbate skin pathology in the context of a weakened skin barrier.

As studies accumulate showing the key role of the native skin microbiome in local immune development and homeostasis, there is a temptation to conclude commensal microbes represent suitable candidates for interventional probiotic therapy. Although we and others (Uberoi et al. 8 2021; Wang et al. 2021) find that depletion of the skin microbiome significantly decreases the ability of the skin to recover from damage, we also observe that supplementation of the native microbiome with additional bioburden of commensals is decidedly detrimental. In developing probiotic therapies, it is thus critical to consider the context of microbial exposure and the range of immune responses that may result therein.

## Data and materials availability

Sequencing data are available as BioProject PRJNA1047182. Other data and code used for analysis and visualization are available on Github: https://github.com/vedomics/commensal_staph.

## Competing interests

Authors declare they have no competing interests

## Acknowledgements

We would like to thank Yiyin Erin Chen for helpful conversations and manuscript review, the Ragon Institute Flow Core (director Michael Waring) for training and assistance in flow cytometry and multiplex ELISA analysis, the MIT Division of Comparative Medicine (DCM) veterinary and husbandry staff, MIT DCM comparative pathology staff member Caroline Atkinson for helpful discussions and sample processing, the Koch Institute Genomics Core and MIT BioMicro Center (director Stuart Levine) for sequencing, and members of the Lieberman lab for useful discussions and comments on the manuscript.

## Funding

Dr. Theron G Randolph Pilot Program administered by the MIT Center for Environmental Health Sciences.

National Institute of Environmental Health Sciences award P30-ES002109 (Koch Institute Core Facility)

## Author contributions

Conceptualisation: TDL, VDK, LM

Methodology: TDL, VDK, LM

Animal work: VDK, LM

Flow cytometry: VDK

Microbiome analysis: VDK

Transcriptomics: LM

Protein analysis: LM

Histopathological analysis: MB

Visualization: VDK

Writing - original draft: VDK, LM

Writing - review and editing: TDL, VDK, LM, MB

## Methods

### Animals

Eight-week-old C57BL/6 male and female mice were purchased from Taconic Biosciences (New York) and housed in individually ventilated cages in a specific pathogen free facility under the Division of Comparative Medicine (DCM) at MIT. Female mice were housed in groups of 3-5 animals per cage and male mice were housed singly to avoid any fighting behaviors that could compromise the skin barrier. Mice were maintained on a 12h light-dark cycle at ambient humidity in sterilized cages and given sterile food and water to limit exposure to facility bacteria. Cages were changed at the beginning of each experiment or every week. All mouse experiments were conducted under protocols approved by MIT DCM IACUC (protocol number 0121-016-24).

### Animal experiments study design

Both male and female mice were used in experiments to establish the tape-stripping model. We did not observe a significant difference in the delayed healing response to commensal bacteria as a result of sex (Fig. 2a and Fig. S1 include both male and female mice across experimental groups). The effect size of increased TEWL and severity score AUC observed as a consequence of Sepi-TDL44-exposure in a pilot experiment (total N=17) was used for power calculations which determined the total sample size per group required for statistical significance (N=12); this number was split between 2-3 cohorts of animals (N=4-5 per cohort) to ensure reproducibility across experimental repeats. Unless noted in text, experimental groups (bacterial exposure or antibiotic treatment) were compared to a control group which undergoes hair removal and tape-stripping damage and has PBS vehicle applied to skin daily after damage. Animals were randomly assigned to experimental conditions.

Animals were excluded from analysis if hair regrowth occurred during the three day interval between hair removal and tape-stripping. Comparable tape-stripping damage was not possible or if hair regrew significantly by 24h post-tape-stripping.

### Statistical analysis

Throughout the text, when comparing an experimental group to the control a two-sided Wilcoxon rank-sum test was used to compare the medians of the two groups. Only PBS control animals from cohorts that included the experimental group of interest were used for comparison. When multiple groups were being compared to the control this was followed by the Benjamini Hochberg correction for multiple hypothesis testing. Pearson’s correlation coefficient was calculated to quantify correlation. Likelihood ratio test followed by Benjamini Hochberg correction was used for differential expression analysis of transcriptomic data. Fisher’s exact test was used to calculate enrichment. Statistical analysis was performed in R 4.1.3 or python (SciPy).

### Tape-stripping Barrier Damage Model

Mice were shaved and depilated (Nair) 72 hours prior to barrier damage. Depilation itself did not induce inflammation or damage to the skin barrier by histopathological assessment (Fig. S1). Tape-stripping was performed by applying and removing a Tegaderm bandage (3M) to the depilated dorsal flank skin 10 times, resulting in significant disruption of the epidermis observed as skin reddening and glistening. Mice were allowed to recover from damage and anesthesia in a heated recovery cage and then placed back in their home cage.

To assess barrier damage, transepidermal water loss (TEWL) was measured using a noninvasive probe (Tewameter, C+K). Continuous TEWL measurements for each animal were taken for up to 20s, or until the standard deviation between readings fell below 0.5 g/m^2^/h. TEWL values immediately following tape-stripping were between 60–80 g/m^2^/h. TEWL was measured daily following damage and severity score was assessed beginning 48h after damage (prior to 48h no morphological changes in skin were apparent). For consistency, the same experimenter performed all tape-stripping and TEWL measurements. Disease severity score assessed gross morphological changes during healing and included skin thickness (0-3), scale (0-3), and erythema (0-3). To limit duration of daily anesthesia required for both bacterial application and TEWL and score assessment, all manipulations were performed by the same experimenter and thus experimenters could not perform blinded scoring. Mice were euthanized at the end of the experiment by 5% C02 inhalation.

### Application of bacteria or vehicle control

100μl of phosphate-buffered saline vehicle (PBS, ThermoFisher) or washed bacterial cultures (10^9^ CFUs) was pipetted onto damaged flank skin and then gently rolled across the skin surface using a sterile swab (Puritan Medical Products).

### Bacterial growth and preparation

Bacteria was subcultured (1:100) from overnight cultures and grown to exponential phase at 37°C with shaking. Bacterial cells were then washed twice with PBS and resuspended at a concentration of 10^10^ CFU/ml except *L. reuteri* which was resuspended at a concentration of 10^9^ CFU/ml. *S. epidermidis* (TDL44 and TDL105) and *S. hominis* strains were isolated from healthy volunteers and are part of the lab strain collection. *S. epidermidis* NIHLM087 was a generous gift from Dr. Chris Voigt. S*. xylosus* was isolated from mouse skin from animals housed in the MIT animal facilities. *S. aureus* USA300 LAC (Diep et al. 2006) was used and was a generous gift from Dr. Isaac Chiu. *C. accolens* (ATCC49725) and *L. reuteri* (ATCC23272) were obtained from the ATCC.

### Bacterial Inactivation

Exponential phase cultures of *S. epidermidis* (TDL44) were washed and resuspended in PBS to a density of 10^10^ CFU/mL. Cells were heat-killed by exposure to 95°C for 1 hour. Ethanol inactivation was achieved by resuspending cells in 80% ethanol (freshly prepared) for 2 hours on ice, with periodic mixing. The ethanol inactivated cells were pelleted and washed with ice-cold PBS twice and resuspended at the original concentration. For both methods, killing was confirmed by plating the inactivated cells on tryptic soy agar.

### Bacterial Enumeration from Murine Skin

Animals were euthanized via C02 inhalation, and a ∼1cm^2^ area of flank skin was collected in PBS kept on ice. Skin sections were weighed before being minced and homogenized using a TissueLyserII (Qiagen). Dilutions of skin homogenate were plated on Mannitol Salt Agar (Oxoid) and colonies enumerated.

### Antibiotic Treatment

Antibiotic depletion was performed as per (Uberoi et al. 8 2021). Briefly, metronidazole (1g/L, Sigma), sulfamethoxazole (0.8g/L, Sigma), trimethoprim (0.16g/L, Sigma), cephalexin (4g/L, Sigma) and Baytril (0.025g/L, Sigma) were dissolved in drinking water containing Splenda (1 packet/250ml) as a sweetener and was provided to mice for two weeks prior to tape-stripping barrier damage and throughout the barrier damage protocol to the endpoint of the experiment. Cages were changed 3 times/week to ensure decreased microbial burden in antibiotic treated mice. Control cages were given drinking water containing Splenda and cages were changed once per week to ensure microbial diversity.

### Tissue dissociation and Flow Cytometry

Flank skin was harvested from euthanized animals and placed in RPMI 1640 with L-glutamine and HEPES (Gibco), containing 10% serum (Gibco). Tissue was minced and digested in RPMI with 1% serum, 0.1mg/mL DNase 0.25mg/mL TL liberase (Roche) overnight at 37°C with 5% C02. The digestion reaction was quenched using 10mL of RPMI 1640 with 10% serum and 1mM EDTA. Digested cells were filtered through a 70μM filter before being washed twice with PBS. After antibody blocking (CD16/32, ebioscience), cells were stained with an amine reactive live/dead dye (efluor506, Thermo Scientific) and an antibody panel (Ly6G-PE, F4/80-BrilliantViolet600, CD11c-BV711, MHCII-Alexa700, CD11b-PECy5, Ly-gC-Alexa488, CD45-efluor450, CD3-APC, CD8-Alexa488, CD4-SuperBright600, Thermofisher) and fixed using CytoFix (BD) for 30m at 4°C in the dark. Fixative was washed, and stained cells were captured on a BD 5L LSR Fortessa.

### DNA Extraction and 16S Sequencing

The dorsal flank skin of mice was sampled using a swab pre-wetted in a TES solution containing 1.2% Triton X-100 (Sigma) vigorously rubbed on the skin surface. The swab was stored in DNA/RNA Shield (Zymo) in a Zymo Bead Bashing Lysis tube (Zymo) and frozen. Fecal pellets were also collected and frozen dry. DNA was extracted from both skin and fecal samples using the ZymoBIOMICS 96 DNA kit (Zymo), following manufacturer’s protocol. The 16S region of the bacterial rRNA gene was amplified using V1-V3 primers (27F - 543R). Libraries were prepared for sequencing following the Hackflex protocol (Gaio et al. 2022). Nextera-compatible primers (IDT) were used for index PCR and amplicons were purified using DNA-binding beads (Cytiva) for size selection.

### Microbiome Analysis

Sequencing was performed at the MIT BioMicro Center on an Illumina Miseq using 300bp paired-ends reads to an average depth of 300,000 reads per sample. All data processing was done using QIIME2 (v2021.2) (Bolyen et al. 2019). Only forward reads were used, as reverse read quality was too low to overlap pairs. Adapters were trimmed using the cutadapt plugin for QIIME2, and data were denoised using DADA2 (Callahan et al. 2016) to generate the amplicon sequence variant (ASV) table. As described previously (Gupta et al. 2023), a custom classifier based on the SILVA database (v132) (Quast et al. 2013) was used for taxonomic assignment. Exported taxa abundance from QIIME2 were analyzed using R version 4.1.3 and phyloseq version 1.38 (McMurdie and Holmes 2013). Taxa unassigned at the phylum level and below, as well as taxa assigned to eukaryotes, were removed. Taxa with fewer than 250 reads across all samples were removed and analyses performed on the remaining subset.

### RNA Isolation and cDNA library synthesis

A small (∼1cm^2^) piece of skin from the area subjected to damage (when applicable) was dissected immediately after euthanasia and either placed in RNAlater (Invitrogen) or snap-frozen in liquid nitrogen. Samples placed in RNAlater were placed on ice for 1-4 hours and then frozen at −80°C. Snap-frozen samples were placed on dry ice and then transferred to −80°C. To extract RNA, samples were placed in a 2ml tube with 1ml of QIAzol (Qiagen) containing a sterile 4.5mm ball bearing and homogenized with 2 rounds of bead-beating (2m30s, 30 beats/s) in a TissueLyserII (Qiagen). The lysate was then transferred to a new tube and 1/5 volume of chloroform (Avantor) added and lysate shaken by hand for 15 seconds. This tube was centrifuged for 15 minutes at 4°C at 12,000xg and the aqueous layer transferred to a new tube and 1 volume of 70% ethanol added. RNA was then purified using a PureLink RNA kit with on-column DNase treatment (Invitrogen) as per manufacturer’s guidelines. RNA was quantified using a Nanodrop and ∼500ng was used for cDNA synthesis as per the Smart-seq2 protocol (Picelli et al. 2014). cDNA libraries were fragmented and prepared for sequencing following the Hackflex protocol (Gaio et al. 2022) modified as follows: bead-linked transposase was diluted 1:5 in buffer and used for tagmentation as per protocol. Nextera-compatible unique dual index primers (IDT) were used for index PCR and fragments were purified using DNA-binding beads (Cytiva) for double-sided size selection.

### RNA sequencing and differential expression analysis

Libraries were sequenced using a Novaseq S4 flow cell (50 basepair single-end reads) at an average sequencing depth of 25 million reads per sample. Reads were pseudoaligned to the mouse transcriptome using kallisto ((Bray et al. 2016), default parameters) and differential expression analysis performed using sleuth ((Pimentel et al. 2017), default parameters). Differential expression analysis included animals from multiple cohorts sacrificed at multiple timepoints after damage. Experimental cohort was included as a variable such that genes shown are differentially expressed between experimental conditions at the timepoint shown regardless of cohort effects.

Statistically significant genes (likelihood ratio test, p-value <0.05, Benjamini-Hochberg correction for multiple comparisons) from each timepoint were used as input for pathway enrichment analysis using clusterProfiler (Wu et al. 2021) and the GO database (release date 2023-01-01, version 10.5281/zenodo.7504797, (Ashburner et al. 2000; Gene Ontology Consortium et al. 2023). Pathway enrichment analysis was performed on the entire set of differentially expressed genes as well as exclusively on the subset of genes annotated as part of the immune response in the Mouse Genome Database (Blake et al. 2021). Enrichment was calculated based on over-representation analysis and a one-sided Fisher’s exact test, with Benjamini-Hochberg correction for multiple comparisons. Pathways were then filtered to include those with at least 5 differentially expressed genes and curated for processes of interest. Uncurated clusterProfiler output for immune genes and non-immune genes is contained in Table S1 and Table S2.

### Histology / pathology

A representative strip of skin taken from the most damaged section of the tapestripped area was fixed in 10% neutral buffered formalin, embedded in paraffin, sectioned at 5μm thickness and stained with hematoxylin and eosin. Sections were scored by a board-certified veterinary pathologist. Skin was assessed for the presence of serocellular crust as follows: 0=none, 1=little/occasionally observed, 2=severe crust formation. The following criteria were used for scoring dermal and subcutaneous neutrophil infiltrates: 0=none, 1=minimal, 2=moderate and 3=marked. The total score per animal is shown (Fig. S1). Slides were imaged using a digital slide scanner (Aperio) at 20x magnification.

The thickness of the top layer of skin (epidermis and/or serocellular crust depending on degree of epidermal ulceration) was quantified using ImageJ and a custom python script as follows: two representative sections per animal were annotated in ImageJ such that x-y coordinates of the top and bottom of the epidermis were saved as a text file. The distance between the top and bottom of that area was then calculated using euclidean distance in python and averaged across both sections per animal.

### Immunoassay for skin cytokine protein quantification

Skin samples were snap-frozen in liquid nitrogen immediately after sacrifice and stored at −80°C. To prepare whole skin lysate, frozen samples were diced using a scalpel then 500ul of lysis buffer (RIPA with 1mM PMSF) was added per 100mg of sample and two sterile 4.5mm ball-bearing were added to the tube prior to mechanical dissociation using the TissueLyserII (Qiagen): 25 beats/s for 3 minutes, repeated up to 3 times until homogenized. Lysate was centrifuged at 4C for 10 minutes at 16,000xg to remove unhomogenized debris and then supernatant was aliquoted and frozen at −80°C. Total protein was measured using a Bradford Assay (BioRad). Skin lysates were diluted 1:10 in ProcartaPlex Universal Assay Buffer prior to use in custom Procartaplex Immunoassay (ThermoFisher) analyzed using the FlexMap3D (Luminex). Skin lysates were subjected to acid-ethanol extraction (Flanders et al. 2016), lyophilized and resuspended in Procartaplex Universal Assay Buffer and stored at −80°C prior to measurement of TGF-B using Procartaplex Single-plex assay (ThermoFisher).

## Notes

### Competing Interest Statement

The authors have declared no competing interest.

## References

1. Acosta EM, Little KA, Bratton BP, Lopez JG, Mao X, Payne AS, et al. Bacterial DNA on the skin surface overrepresents the viable skin microbiome. Elife [Internet]. 2023 Jun 30;12. Available from: 10.7554/eLife.87192

2. Adair-Kirk TL, Senior RM. Fragments of extracellular matrix as mediators of inflammation. Int J Biochem Cell Biol. 2008;40(6-7):1101–10.

3. Asarch A, Barak O, Loo DS, Gottlieb AB. Th17 cells: a new paradigm for cutaneous inflammation. J Dermatolog Treat. 2008;19(5):259–66.

4. Ashburner M, Ball CA, Blake JA, Botstein D, Butler H, Cherry JM, et al. Gene ontology: tool for the unification of biology. The Gene Ontology Consortium. Nat Genet. 2000 May;25(1):25–9.

5. Belheouane M, Vallier M, Čepić A, Chung CJ, Ibrahim S, Baines JF. Assessing similarities and disparities in the skin microbiota between wild and laboratory populations of house mice. ISME J. 2020 Oct;14(10):2367–80.

6. Bergmeier V, Etich J, Pitzler L, Frie C, Koch M, Fischer M, et al. Identification of a myofibroblast-specific expression signature in skin wounds. Matrix Biol. 2018 Jan;65:59–74.

7. Blake JA, Baldarelli R, Kadin JA, Richardson JE, Smith CL, Bult CJ, et al. Mouse Genome Database (MGD): Knowledgebase for mouse-human comparative biology. Nucleic Acids Res. 2021 Jan 8;49(D1):D981–7.

8. Bolyen E, Rideout JR, Dillon MR, Bokulich NA, Abnet CC, Al-Ghalith GA, et al. Reproducible, interactive, scalable and extensible microbiome data science using QIIME 2. Nat Biotechnol. 2019 Aug;37(8):852–7.

9. Bray NL, Pimentel H, Melsted P, Pachter L. Near-optimal probabilistic RNA-seq quantification. Nat Biotechnol. 2016 May;34(5):525–7.

10. Burian M, Bitschar K, Dylus B, Peschel A, Schittek B. The Protective Effect of Microbiota on S. aureus Skin Colonization Depends on the Integrity of the Epithelial Barrier. J Invest Dermatol. 2017 Apr;137(4):976–9.

11. Byrd AL, Deming C, Cassidy SKB, Harrison OJ, Ng WI, Conlan S, et al. and strain diversity underlying pediatric atopic dermatitis. Sci Transl Med [Internet]. 2017 Jul 5;9(397). Available from: 10.1126/scitranslmed.aal4651

12. Cai Y, Shen X, Ding C, Qi C, Li K, Li X, et al. Pivotal role of dermal IL-17-producing γδ T cells in skin inflammation. Immunity. 2011 Oct 28;35(4):596–610.

13. Callahan BJ, McMurdie PJ, Rosen MJ, Han AW, Johnson AJA, Holmes SP. DADA2: High-resolution sample inference from Illumina amplicon data. Nat Methods. 2016 Jul;13(7):581–3.

14. Canovas J, Baldry M, Bojer MS, Andersen PS, Gless BH, Grzeskowiak PK, et al. Cross-Talk between Staphylococcus aureus and Other Staphylococcal Species via the agr Quorum Sensing System. Front Microbiol [Internet]. 2016 [cited 2020 Dec 3];7. Available from: 10.3389/fmicb.2016.01733/full

15. Cau L, Williams MR, Butcher AM, Nakatsuji T, Kavanaugh JS, Cheng JY, et al. Staphylococcus epidermidis protease EcpA can be a deleterious component of the skin microbiome in atopic dermatitis. J Allergy Clin Immunol. 2021 Mar;147(3):955–66.e16.

16. Chang HW, Yan D, Singh R, Liu J, Lu X, Ucmak D, et al. Alteration of the cutaneous microbiome in psoriasis and potential role in Th17 polarization. Microbiome. 2018 Sep 5;6(1):154.

17. Clausen ML, Agner T, Lilje B, Edslev SM, Johannesen TB, Andersen PS. Association of Disease Severity With Skin Microbiome and Filaggrin Gene Mutations in Adult Atopic Dermatitis. JAMA Dermatol. 2018 Mar 1;154(3):293–300.

18. Conwill A, Kuan AC, Damerla R, Poret AJ, Baker JS, Tripp AD, et al. Anatomy promotes neutral coexistence of strains in the human skin microbiome. Cell Host Microbe. 2022 Feb 9;30(2):171–82.e7.

19. Diep BA, Gill SR, Chang RF, Phan TH, Chen JH, Davidson MG, et al. Complete genome sequence of USA300, an epidemic clone of community-acquired meticillin-resistant Staphylococcus aureus. Lancet. 2006 Mar 4;367(9512):731–9.

20. Flanders KC, Yang YA, Herrmann M, Chen J, Mendoza N, Mirza AM, et al. Quantitation of TGF-β proteins in mouse tissues shows reciprocal changes in TGF-β1 and TGF-β3 in normal vs neoplastic mammary epithelium. Oncotarget. 2016 Jun 21;7(25):38164–79.

21. Gaio D, Anantanawat K, To J, Liu M, Monahan L, Darling AE. Hackflex: low-cost, high-throughput, Illumina Nextera Flex library construction. Microb Genom. 2022 Jan;8(1):000744.

22. Gene Ontology Consortium, Aleksander SA, Balhoff J, Carbon S, Cherry JM, Drabkin HJ, et al. The Gene Ontology knowledgebase in 2023. Genetics [Internet]. 2023 May 4;224(1). Available from: 10.1093/genetics/iyad031

23. Gevers D, Kugathasan S, Denson LA, Vázquez-Baeza Y, Van Treuren W, Ren B, et al. The treatment-naive microbiome in new-onset Crohn’s disease. Cell Host Microbe. 2014 Mar 12;15(3):382–92.

24. Gimblet C, Meisel JS, Loesche MA, Cole SD, Horwinski J, Novais FO, et al. Cutaneous Leishmaniasis Induces a Transmissible Dysbiotic Skin Microbiota that Promotes Skin Inflammation. Cell Host Microbe. 7 2017;22:13–24.e4.

25. Grice EA, Kong HH, Conlan S, Deming CB, Davis J, Young AC, et al. Topographical and temporal diversity of the human skin microbiome. Science. 2009 May 29;324(5931):1190–2.

26. Gupta S, Poret AJ, Hashemi D, Eseonu A, Yu SH, D’Gama J, et al. Cutaneous Surgical Wounds Have Distinct Microbiomes from Intact Skin. Microbiol Spectr. 2023 Feb 14;11(1):e0330022.

27. Hattori N, Mochizuki S, Kishi K, Nakajima T, Takaishi H, D’Armiento J, et al. MMP-13 plays a role in keratinocyte migration, angiogenesis, and contraction in mouse skin wound healing. Am J Pathol. 2009 Aug;175(2):533–46.

28. Herrick S, Ashcroft G, Ireland G, Horan M, McCollum C, Ferguson M. Up-regulation of elastase in acute wounds of healthy aged humans and chronic venous leg ulcers are associated with matrix degradation. Lab Invest. 1997 Sep;77(3):281–8.

29. Joshi N, Pohlmeier L, Greenwald MBY, Haertel E, Hiebert P, Kopf M, et al. Comprehensive characterization of myeloid cells during wound healing in healthy and healing-impaired diabetic mice. Eur J Immunol. 9 2020;50:1335–49.

30. Khadka VD, Key FM, Romo-González C, Martínez-Gayosso A, Campos-Cabrera BL, Gerónimo-Gallegos A, et al. The Skin Microbiome of Patients With Atopic Dermatitis Normalizes Gradually During Treatment. Front Cell Infect Microbiol. 2021 Sep 24;11:720674.

31. Komatsu N, Saijoh K, Kuk C, Liu AC, Khan S, Shirasaki F, et al. Human tissue kallikrein expression in the stratum corneum and serum of atopic dermatitis patients. Exp Dermatol. 2007 Jun;16(6):513–9.

32. Kong HH, Oh J, Deming C, Conlan S, Grice EA, Beatson MA, et al. Temporal shifts in the skin microbiome associated with disease flares and treatment in children with atopic dermatitis. Genome Res. 2012 May;22(5):850–9.

33. Landén NX, Li D, Ståhle M. Transition from inflammation to proliferation: a critical step during wound healing. Cell Mol Life Sci. 2016 Oct 1;73(20):3861–85.

34. Lauer G, Sollberg S, Cole M, Flamme I, Stürzebecher J, Mann K, et al. Expression and proteolysis of vascular endothelial growth factor is increased in chronic wounds. J Invest Dermatol. 2000 Jul;115(1):12–8.

35. Lecron JC, Charreau S, Jégou JF, Salhi N, Petit-Paris I, Guignouard E, et al. IL-17 and IL-22 are pivotal cytokines to delay wound healing of infected skin. Front Immunol. 2022 Oct 7;13:984016.

36. Leech JM, Dhariwala MO, Lowe MM, Chu K, Merana GR, Cornuot C, et al. Toxin-Triggered Interleukin-1 Receptor Signaling Enables Early-Life Discrimination of Pathogenic versus Commensal Skin Bacteria. Cell Host Microbe. 2019 Dec 11;26(6):795–809.e5.

37. Lee J, Rodero MP, Patel J, Moi D, Mazzieri R, Khosrotehrani K. Interleukin-23 regulates interleukin-17 expression in wounds, and its inhibition accelerates diabetic wound healing through the alteration of macrophage polarization. FASEB J. 2018 Apr;32(4):2086–94.

38. Lin AM, Rubin CJ, Khandpur R, Wang JY, Riblett M, Yalavarthi S, et al. Mast cells and neutrophils release IL-17 through extracellular trap formation in psoriasis. J Immunol. 2011 Jul 1;187(1):490–500.

39. Linehan JL, Harrison OJ, Han SJ, Byrd AL, Vujkovic-Cvijin I, Villarino AV, et al. Non-classical Immunity Controls Microbiota Impact on Skin Immunity and Tissue Repair. Cell. 2018 Feb 8;172(4):784–96.e18.

40. McMurdie PJ, Holmes S. phyloseq: an R package for reproducible interactive analysis and graphics of microbiome census data. PLoS One. 2013 Apr 22;8(4):e61217.

41. Michopoulou A, Rousselle P. How do epidermal matrix metalloproteinases support re-epithelialization during skin healing? Eur J Dermatol. 2015 Apr;25 Suppl 1:33–42.

42. Naik S, Bouladoux N, Linehan JL, Han SJ, Harrison OJ, Wilhelm C, et al. Commensal-dendritic-cell interaction specifies a unique protective skin immune signature. Nature. 2015 Apr 2;520(7545):104–8.

43. Naik S, Bouladoux N, Wilhelm C, Molloy MJ, Salcedo R, Kastenmuller W, et al. Compartmentalized control of skin immunity by resident commensals. Science. 2012 Aug 31;337(6098):1115–9.

44. Nakagawa S, Matsumoto M, Katayama Y, Oguma R, Wakabayashi S, Nygaard T, et al. Staphylococcus aureus Virulent PSMα Peptides Induce Keratinocyte Alarmin Release to Orchestrate IL-17-Dependent Skin Inflammation. Cell Host Microbe. 2017 Nov 8;22(5):667–77.e5.

45. Nakatsuji T, Chen TH, Narala S, Chun KA, Two AM, Yun T, et al. Antimicrobials from human skin commensal bacteria protect against *Staphylococcus aureus* and are deficient in atopic dermatitis. Sci Transl Med [Internet]. 2017 Feb 22;9(378).Available from: 10.1126/scitranslmed.aah4680

46. Nakatsuji T, Hata TR, Tong Y, Cheng JY, Shafiq F, Butcher AM, et al. Development of a human skin commensal microbe for bacteriotherapy of atopic dermatitis and use in a phase 1 randomized clinical trial. Nat Med. 2021 Apr;27(4):700–9.

47. Nomura H, Suganuma M, Takeichi T, Kono M, Isokane Y, Sunagawa K, et al. Multifaceted Analyses of Epidermal Serine Protease Activity in Patients with Atopic Dermatitis. Int J Mol Sci [Internet]. 2020 Jan 30;21(3). Available from: 10.3390/ijms21030913

48. Paharik AE, Parlet CP, Chung N, Todd DA, Rodriguez EI, Van Dyke MJ, et al. Coagulase-Negative Staphylococcal Strain Prevents Staphylococcus aureus Colonization and Skin Infection by Blocking Quorum Sensing. Cell Host Microbe. 2017 Dec 13;22(6):746–56.e5.

49. Picelli S, Faridani OR, Björklund AK, Winberg G, Sagasser S, Sandberg R. Full-length RNA-seq from single cells using Smart-seq2. Nat Protoc. 2014 Jan;9(1):171–81.

50. Pimentel H, Bray NL, Puente S, Melsted P, Pachter L. Differential analysis of RNA-seq incorporating quantification uncertainty. Nat Methods. 2017 Jul;14(7):687–90.

51. Quast C, Pruesse E, Yilmaz P, Gerken J, Schweer T, Yarza P, et al. The SILVA ribosomal RNA gene database project: improved data processing and web-based tools. Nucleic Acids Res. 2013 Jan;41(Database issue):D590–6.

52. Ramsey MM, Freire MO, Gabrilska RA, Rumbaugh KP, Lemon KP. Staphylococcus aureus Shifts toward Commensalism in Response to Corynebacterium Species. Front Microbiol [Internet]. 2016 [cited 2021 Jan 13];7. Available from: 10.3389/fmicb.2016.01230/full

53. Reshamwala K, Cheung GYC, Hsieh RC, Liu R, Joo HS, Zheng Y, et al. Identification and characterization of the pathogenic potential of phenol-soluble modulin toxins in the mouse commensal Staphylococcus xylosus. Front Immunol. 2022;13:999201.

54. Saleh F, Kheirandish F, Azizi H, Azizi M, Departement, Departement, et al. Molecular diagnosis and characterization of Bacillus subtilis isolated from burn wound in Iran. Res Mol Med. 2014 May 1;2(2):40–4.

55. Singh S, Young A, McNaught CE. The physiology of wound healing. Surgery. 2017 Sep 1;35(9):473–7.

56. Taddese R, Belzer C, Aalvink S, de Jonge MI, Nagtegaal ID, Dutilh BE, et al. Production of inactivated gram-positive and gram-negative species with preserved cellular morphology and integrity. J Microbiol Methods. 2021 May 1;184:106208.

57. Takagi N, Kawakami K, Kanno E, Tanno H, Takeda A, Ishii K, et al. IL-17A promotes neutrophilic inflammation and disturbs acute wound healing in skin. Exp Dermatol. 2 2017;26:137–44.

58. Uberoi A, Bartow-McKenney C, Zheng Q, Flowers L, Campbell A, Knight SAB, et al. Commensal microbiota regulates skin barrier function and repair via signaling through the aryl hydrocarbon receptor. Cell Host Microbe. 8 2021;29:1235–48.e8.

59. Voegeli R, Rawlings AV, Breternitz M, Doppler S, Schreier T, Fluhr JW. Increased stratum corneum serine protease activity in acute eczematous atopic skin. Br J Dermatol. 2009 Jul;161(1):70–7.

60. Wang G, Sweren E, Liu H, Wier E, Alphonse MP, Chen R, et al. Bacteria induce skin regeneration via IL-1β signaling. Cell Host Microbe. 2021 May 12;29(5):777–91.e6.

61. Wanke I, Skabytska Y, Kraft B, Peschel A, Biedermann T, Schittek B. Staphylococcus aureus skin colonization is promoted by barrier disruption and leads to local inflammation. Exp Dermatol. 2013;22(2):153–5.

62. Wei M, Flowers L, Knight SAB, Zheng Q, Murga-Garrido S, Uberoi A, et al. Harnessing diversity and antagonism within the pig skin microbiota to identify novel mediators of colonization resistance to methicillin-resistant Staphylococcus aureus. mSphere. 2023 Aug 24;8(4):e0017723.

63. Williams MR, Cau L, Wang Y, Kaul D, Sanford JA, Zaramela LS, et al. Interplay of Staphylococcal and Host Proteases Promotes Skin Barrier Disruption in Netherton Syndrome. Cell Rep. 2020 Mar 3;30(9):2923–33.e7.

64. Williams MR, Costa SK, Zaramela LS, Khalil S, Todd DA, Winter HL, et al. Quorum sensing between bacterial species on the skin protects against epidermal injury in atopic dermatitis. Sci Transl Med [Internet]. 5 2019;11. Available from: 10.1126/scitranslmed.aat8329

65. Won YS, Kwon HJ, Oh GT, Kim BH, Lee CH, Park YH, et al. Identification of Staphylococcus xylosus isolated from C57BL/6J-Nos2(tm1Lau) mice with dermatitis. Microbiol Immunol. 2002;46(9):629–32.

66. Wu T, Hu E, Xu S, Chen M, Guo P, Dai Z, et al. clusterProfiler 4.0: A universal enrichment tool for interpreting omics data. Innovation (Camb). 2021 Aug 28;2(3):100141.

67. Zheng Y, Hunt RL, Villaruz AE, Fisher EL, Liu R, Liu Q, et al. Commensal Staphylococcus epidermidis contributes to skin barrier homeostasis by generating protective ceramides. Cell Host Microbe. 2022 Mar 9;30(3):301–13.e9.

68. Zomer HD, Trentin AG. Skin wound healing in humans and mice: Challenges in translational research. J Dermatol Sci. 2018 Apr 1;90(1):3–12.

