## Supplemental Figures for "Commensal skin bacteria exacerbate inflammation and delay skin healing"

### Supplemental Data

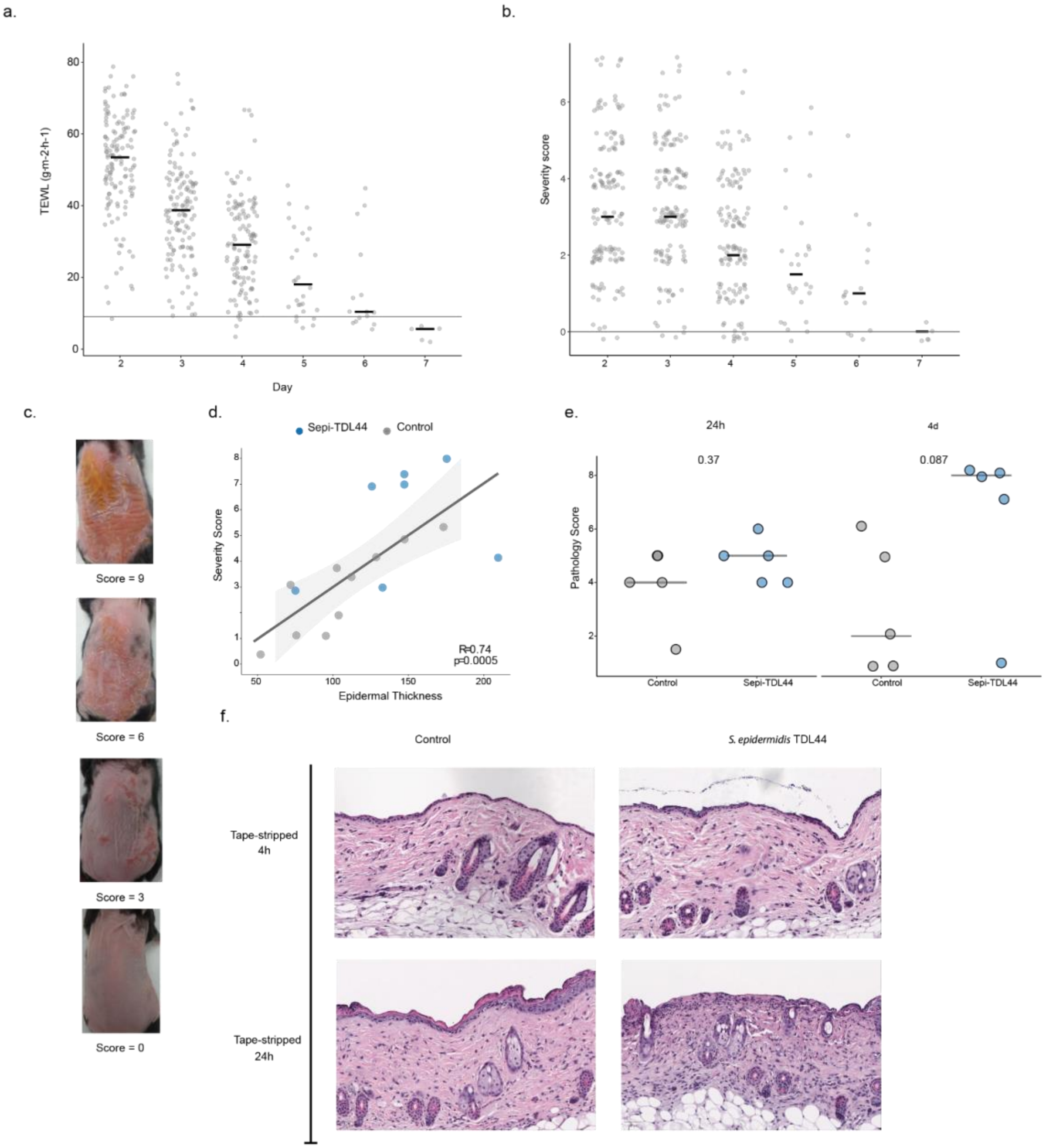


**Supplemental Figure 1: *S. epidermidis* induced inflammation and delayed barrier repair at multiple timepoints.**

Adult mice had hair removed three days prior to tape-stripping damage (day 0). a and b) Immediately following damage and daily until the endpoint, PBS was applied to skin and TEWL was measured and severity score assessed for up to 7 days. a) TEWL was elevated 2 days after damage and returned to baseline level measured on intact skin (average baseline shown as horizontal line) by 7 days after damage. b) Mouse skin thickness (forceps measurement), scale (area) and erythema (redness) were scored for each mouse from 0-3. Cumulative severity score is shown for days 2 through 7 after damage. Skin was comparable to healthy skin (score of 0) by

7 days after damage. Values have been jittered to better represent overlapping data points. c) Representative images of mice with skin severity scores as indicated beneath each picture. d-f) PBS or Sepi-TDL44 was applied to skin immediately after tape-stripping and then daily until experimental endpoint. d) Raw skin severity score correlates significantly with epidermal thickness measured from histology sections (H&E-stained sections of a subset of animals were imaged and average epidermal thickness calculated using ImageJ). Pearson's correlation coefficient R=0.74, P=0.0005, control N=11, Sepi-TDL44 N=6. e) H&E-stained sections were examined by a veterinary pathologist and scored for crust and neutrophil infiltrates of dermis and subcutaneous fat. Cumulative pathology scores are shown and were increased in Sepi-TDL44 4-exposed mice 4 days after tape-stripping damage (P=0.087, control N=5, Sepi-TDL44 N=5 per timepoint). f) Representative H&E-stained sections from animals sacrificed 4h (middle) and 24h (bottom) after tape-stripping damage and application of either vehicle or Sepi-TDL44 . By 24h, there was clear regeneration of the epidermis in control mice, absent from Sepi-TDL44 mice. Instead, Sepi-TDL44-exposed mice showed neutrophil dominant inflammation and epidermal ulceration. Throughout, symbols indicate individual mice. Bars in a, b and e indicate median. Rank sum test was used to compare groups.

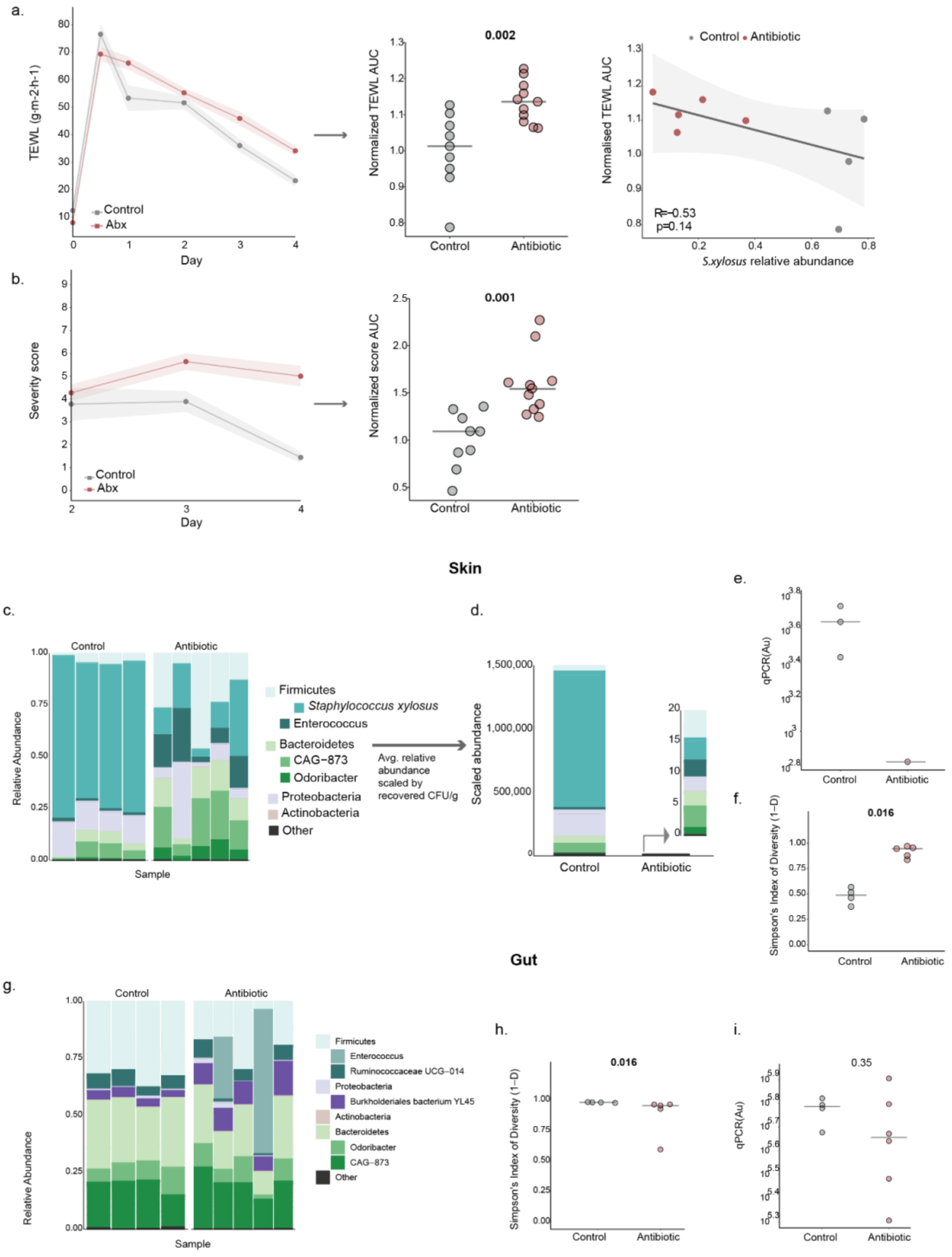


**Supplemental Figure 2: Oral antibiotic treatment depleted the skin microbiome with minimal impact on gut microbiome.**

Mice were given an oral antibiotic cocktail designed to target the skin microbiome for two weeks prior to tape-stripping barrier damage. a) Daily TEWL values (left) were summarized per animal using AUC and normalized to cohort control (no antibiotic) average (middle) and correlated with relative abundance of *S. xylosus* as measured by 16s sequencing (right). b) Daily severity score measurements (left) were summarized per animal using AUC and normalized to cohort control (right). c-i) Animals were sacrificed 4 days after barrier damage and skin was swabbed to collect skin microbiome samples; colon samples were collected for gut microbiome analysis (control N=4 and antibiotics N=5). DNA was extracted and 16s amplicon sequencing performed to define the microbiome. c-f) Analysis of the skin microbiome. There was a dramatic change in skin microbiome composition as a result of antibiotic treatment. c) Relative abundance of bacterial taxa at multiple phylogenetic levels is shown. Bars indicate individual mice. *S. xylosus* dominates control mouse skin microbiome while many taxa comprise skin microbiome of antibiotic-treated mice. d) To demonstrate the depletion of the skin microbiota in antibiotic-treated mice, scaled abundance was calculated and is shown here. Scaled abundance reflects the average relative abundance of each taxa (per experimental group) as measured by sequencing multiplied by the average CFU/g recovered from endpoint skin homogenate CFU (plated on mannitol salt agar). e) Many skin swab samples from antibiotic-treated mice were comparable to water control and thus are not shown; one sample with detectable DNA was significantly lower than skin swab samples collected from control animals consistent with CFU plating (Fig. 1b). f) Simpson's diversity was higher in antibiotic-treated mice compared to controls (P=0.016). g-i) Analysis of the colon microbiome. In most mice analyzed, there were no significant changes in gut microbiome composition as a result of antibiotic administration. g) Relative abundance of bacterial taxa at multiple phylogenetic levels is shown. Bars indicate individual mice. h) Simpson's diversity was significantly decreased (P=0.016) in antibiotic-treated mice, due to one mouse with a dramatic decrease in overall microbiome diversity. i) qPCR with universal 16s primers was used to quantify the total amount of bacterial DNA present in sequenced samples. There was no significant difference in total bacterial DNA detected in control versus antibiotic treated colon samples (P=0.35). For strip-plots, symbols represent mice and bars indicate medians. For line-plots, symbols indicate mean and shading indicates SEM. Rank sum test was used to compare medians.

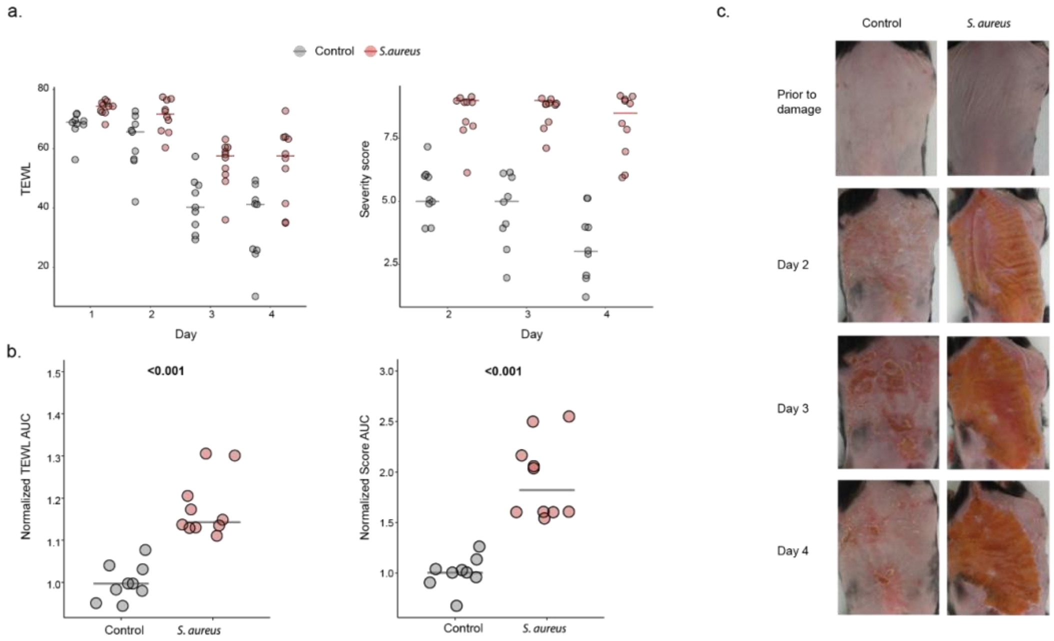


**Supplemental Figure 3: Mice exposed to *S. aureus* after tape-stripping do not heal 4 days post-damage.**

PBS or *S. aureus* USA300 was applied to skin immediately after tape-stripping and then daily until experimental endpoint. control N=9 and *S. aureus* N=10. a) Raw TEWL and severity score values were elevated in *S. aureus*-exposed mice for the duration of the experiment. Raw score is shown to demonstrate the number of *S. aureus*-exposed animals with maximal severity score. b) Normalized TEWL and severity score AUC are significantly higher in *S. aureus*-exposed mice, rank sum test, P=0.00002 and P=0.00002. c) Representative images of control (left) and *S. aureus*-exposed (right) skin prior to damage (top) and throughout the post-damage exposure period. Mice were equivalent prior to tape-stripping and exposure. A thick, red crust covered the majority of the tape-stripped skin of *S. aureus*-exposed mice from day two onward compared to the minimal crust and redness observed in PBS control mice.

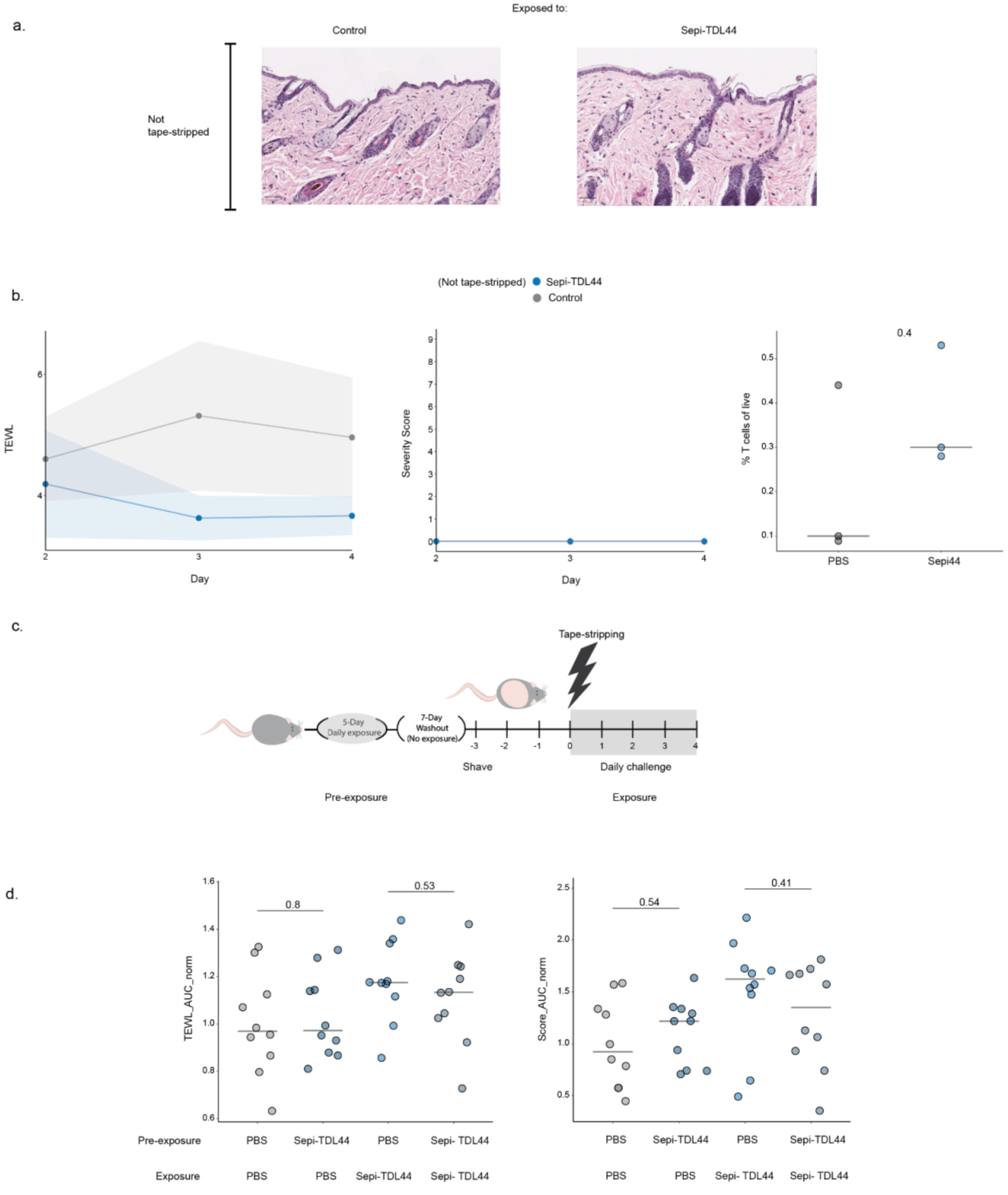


**Supplemental figure 4: Application of *S. epidermidis* to intact skin had no effect on skin barrier function or subsequent response to tape-stripping damage.**

a and b) Three days after hair removal, either PBS or Sepi-TDL44 was applied to intact skin of mice for four days. A) Representative H&E stained sections after four days of PBS or Sepi-TDL44 application (no damage). Histopathological analysis indicated no change in skin pathology or morphology as a result of Sepi-TDL44 application. b) TEWL and severity score were assessed daily during the exposure period (left). TEWL values remained in the undamaged baseline range (2-10 $g/m^2/h$) and no changes in gross morphology (thickness, erythema or scale) were noted so all mice had a score of 0 throughout the exposure period. Bulk T-cells (right) were increased in mice exposed to Sepi-TDL44 after seven days of bacterial application on intact skin. c and d) PBS or $10^9$ CFUs of Sepi-TDL44 were gently pipetted and then spread across intact skin (no hair removal) daily for five days followed by a seven-day washout period. Hair was then removed and three days later mice were subjected to one round of tape-stripping damage. Either PBS or Sepi-TDL44 was applied to skin immediately following tape-stripping and then daily until endpoint. Pre-exposure::challenge exposure: PBS::PBS N=11, PBS::Sepi-TDL44 N=10, Sepi-TDL44::PBS N=11, Sepi-TDL44::Sepi-TDL44 N=11. TEWL AUC (e) and severity score AUC (f) normalized to the cohort control (PBS::PBS) was elevated in mice exposed to Sepi-TDL44 during damage compared to mice exposed to PBS during damage regardless of prior exposure. There was no effect of pre-exposure to Sepi-TDL44 when comparing groups exposed to PBS after damage, suggesting that exposure during health to *S. epidermidis* does not improve healing from tape-stripping barrier damage. Symbols indicate individual mice and lines indicate median.

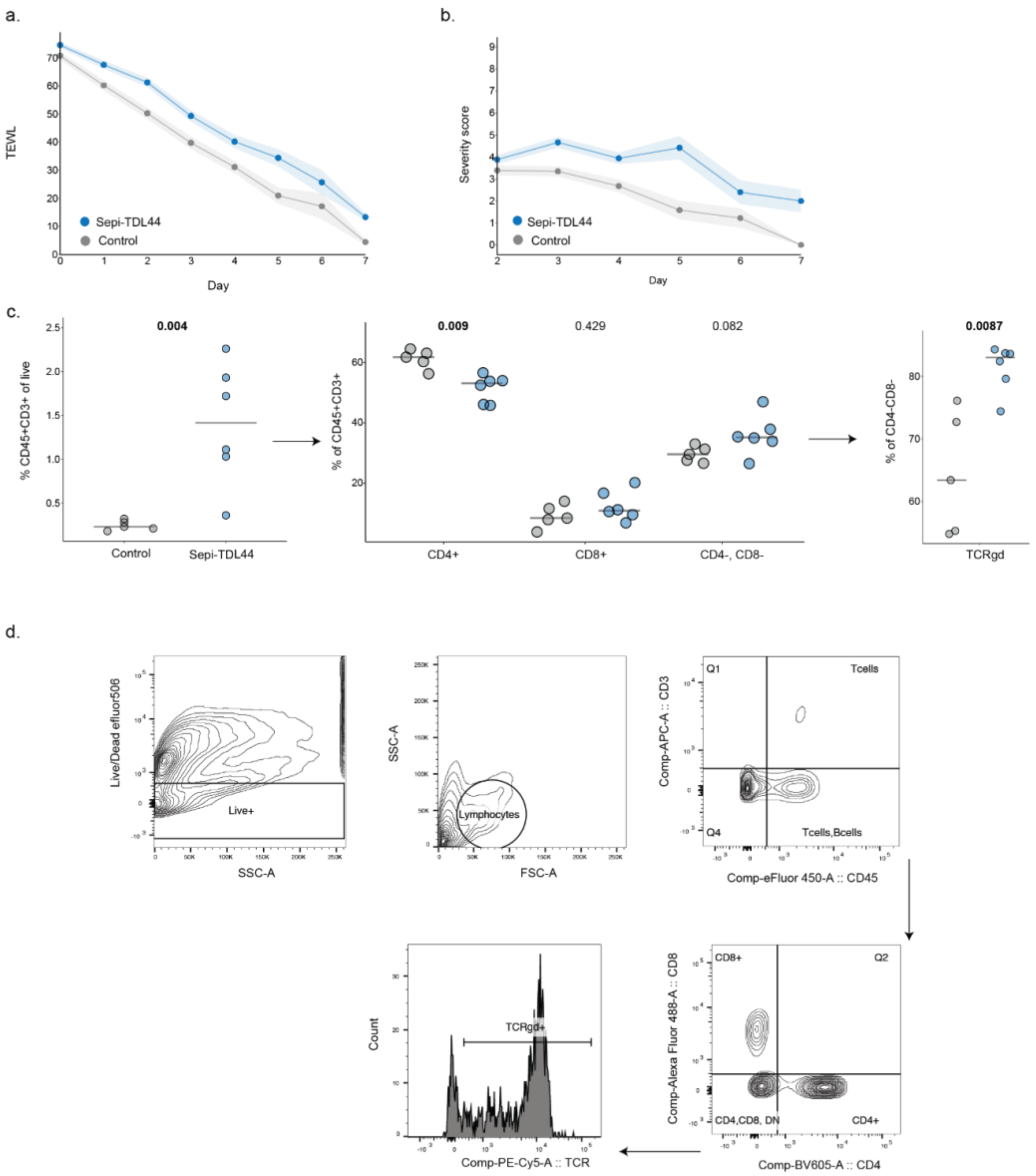


**Supplemental Figure 5: *S. epidermidis* applied after damage induces T cell response.**

Mice had hair removed three days prior to tapestripping damage. PBS or Sepi-TDL44 was applied to skin immediately after tape-stripping and then daily until experimental endpoint (control N=4, Sepi-TDL44 N=6). TEWL was measured daily while severity score was measured starting 2 days after damage (prior to 48h skin is unchanged in gross morphology). a) TEWL and b) severity score were elevated in Sepi-TDL44-exposed mice compared to controls for 7 days after damage. Controls were indistinguishable from un-tape-stripped mice on day seven (score =0) while Sepi-TDL44-exposed mice had non-zero scores on day seven. Symbols indicate mean and shading indicates SEM. c) Flow cytometry was used to measure T cell populations in the skin 7 days after barrier damage. Mice exposed daily to Sepi-TDL44 showed a significant increase in total T-cell abundance (left, P=0.004), with a decrease in CD4+ T cells (middle,

P=0.009) and an increase in γδ T cells (right, P=0.0087). Despite this increase in T cells, mice exposed to Sepi-TDL44 remained more damaged than controls on day seven indicating that T cell induction did not improve barrier function. Symbols indicate individual mice and lines indicate median. d) Gating strategy used to identify T cell subsets. Antibodies used as follows: blocking antibody CD16/32, amine reactive live/dead efluor 506, CD4 SuperBright600, CD8 Alexa488, CD45-PE, CD3-APC, TCRg/d-PeCy5 (Thermofisher).

a.

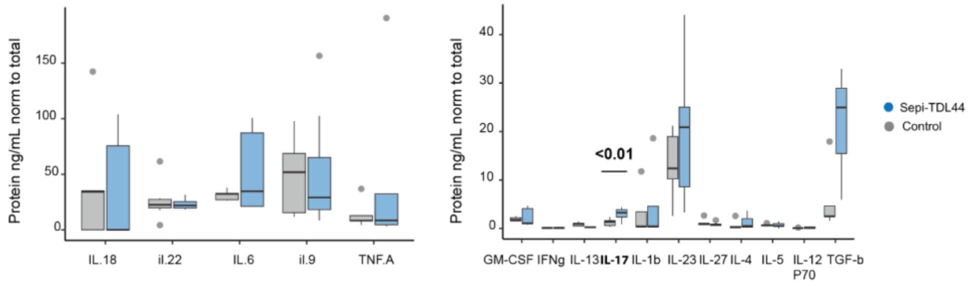


**Supplemental Figure 6:** ***S. epidermidis*** **increased IL-17A protein in skin.**
PBS or Sepi-TDL44 was applied to skin immediately after tape-stripping and then daily until experimental endpoint. Mice were sacrificed on day 3 after damage. Protein was extracted from skin lysates. Total protein was measured by Bradford assay and cytokine proteins were measured using multiplex ELISA. Sepi-TDL44-exposed mice had significantly increased IL-17 protein (rank sum test, P=0.0008). Other increased cytokine protein levels in Sepi-TDL44-exposed mice were not significant after Benjamini Hochberg correction for multiple comparisons. control N=9, Sepi-TDL44 N=11.

a. b. c.

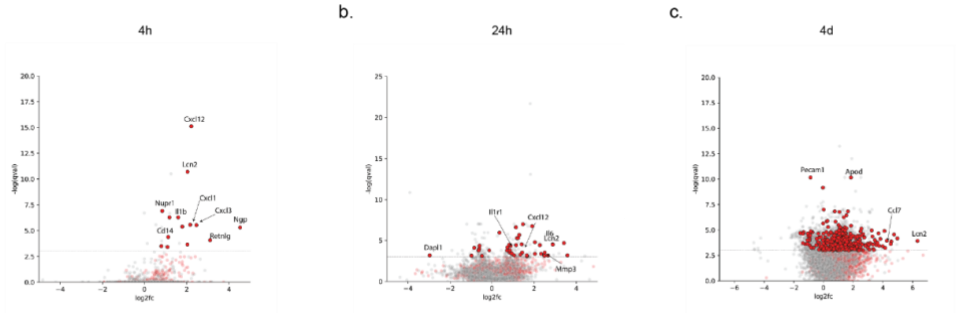


**Supplemental Figure 7:** ***S. epidermidis*** **increased immune gene expression at all timepoints.**
Animals were sacrificed 4h (A), 24h (B) and 4 days (C) after damage and skin samples were used for RNAseq analysis (4h control N=5, Sepi-TDL44 N=5; 24h control N=4, Sepi-TDL44 N=5; 4

days control N=6, Sepi-TDL44 N=9). Transcripts were aligned to the mouse transcriptome and differential expression analysis was performed at the gene-level. Genes annotated as part of the immune response in the Mouse Genome Database are shown in red; all other genes are shown in gray. Dashed line indicates statistical significance of P=0.05. Genes of interest are highlighted with text and arrows.

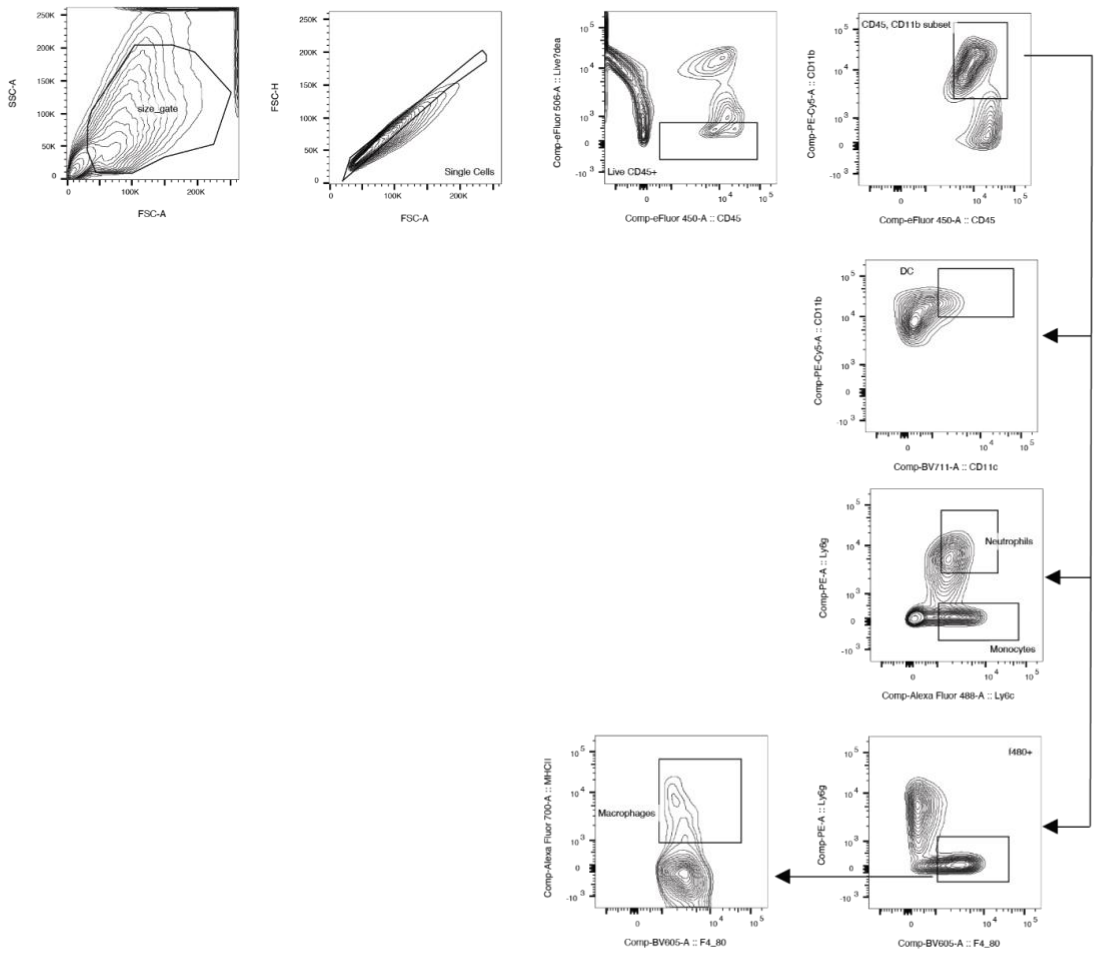


**Supplemental Figure 8: Flow cytometry gating strategy for innate immune cell populations in skin**

Flow cytometry gating strategy to identify innate immune cell subpopulations (neutrophils, macrophages, and conventional dendritic cells) from digested flank skin. Antibodies used were as follows: Ly6G-PE, F4/80-BrilliantViolet600, CD11c-BV711, MHCII-Alexa700, CD11b-PECy5, Ly-gC-Alexa488, CD45-efluor450, CD3-APC.
